# Accelerating the continuous community sharing of digital neuromorphology data

**DOI:** 10.1101/2024.03.15.585306

**Authors:** Carolina Tecuatl, Bengt Ljungquist, Giorgio A. Ascoli

## Abstract

The tree-like morphology of neurons and glia is a key cellular determinant of circuit connectivity and metabolic function in the nervous system of essentially all animals. To elucidate the contribution of specific cell types to both physiological and pathological brain states, it is important to access detailed neuroanatomy data for quantitative analysis and computational modeling. NeuroMorpho.Org is the largest online collection of freely available digital neural reconstructions and related metadata and is continuously updated with new uploads. Earlier in the project, we released multiple datasets together yearly, but this process caused an average delay of several months in making the data public. Moreover, in the past 5 years, >80% of invited authors agreed to share their data with the community via NeuroMorpho.Org, up from <20% in the first 5 years of the project. In the same period, the average number of reconstructions per publication increased 600%, creating the need for automatic processing to release more reconstructions in less time. The progressive automation of our pipeline enabled the transition to agile releases of individual datasets as soon as they are ready. The overall time from data identification to public sharing decreased by 63.7%; 78% of the datasets are now released in less than 3 months with an average workflow duration below 40 days. Furthermore, the mean processing time per reconstruction dropped from 3 hours to 2 minutes. With these continuous improvements, NeuroMorpho.Org strives to forge a positive culture of open data. Most importantly, the new, original research enabled through reuse of datasets across the world has a multiplicative effect on science discovery, benefiting both authors and users.

## Introduction

Brain cells communicate through tree-shaped extensions that process and transmit information^1^. These miniaturized arbors mediate all cognition and behavior^2,a^, and their branching is altered in most neurological and psychiatric conditions^3^, from Alzheimer’s disease^4,5^ to autism spectrum disorders^6–8^. Scientists use microscopes to trace these structures into computer files allowing investigation of neural function, but the process is slow and costly^9,10^. To enable sharing and reuse of these hard-won data, we created NeuroMorpho.Org, a centrally curated public repository of digitally reconstructed neurons and glia associated with peer-reviewed publications^11^, and corresponding metadata annotation^12,13^. Over 1000 laboratories worldwide contributed more than 260,000 three-dimensional neural tracings that can be browsed by species, brain regions, and cell types^14^. These data are re-used for many scientific applications, including comparative analyses, computational modeling, connectivity quantification, machine learning^15^, advanced visualization, and cellular classification, leading to new discoveries every week.

With data sharing becoming increasingly common^16^, the methods by which sharing takes place are receiving increased attention^17^, including data access and configuration management, as well as how releasing data together with software tools for analysis^18^. It is imperative to facilitate increasingly automated machine access for computational use of data resources together with capabilities of direct user-friendly human interaction for discovery and analysis^19^. Advances in microscopic imaging, labeling techniques for specific cell types, and automatic tracing methods^20^ have resulted in a higher yield of reconstructions per animal^21^. Moreover, improved disposition towards data sharing and larger data sets in the neuroscience community created the need for automatic processing to release more reconstructions in less time^22,23^.

Here we describe the new operational workflow of NeuroMorpho.Org encompassing literature mining, data processing, metadata annotation, and database ingestion for public release. We have progressively automated this pipeline and modernized the database to enable agile releases of individual datasets as soon as they are ready, instead of releasing all datasets together once or twice a year. This improvement was facilitated by the development of backend microservices, running as Docker containers^24^, and of routines for rapid and automated formatting, integrity control, ingestion, and incremental sharing of reconstructions. Additionally, we have developed a wide range of tools for further analysis and discovery^25^, including metadata annotation, search functionality, and data conversion^26–28^. These advancements have increased both the accessibility and usability of morphological tracings.

NeuroMorpho.Org always makes data and metadata freely available for download, with no registration required, resulting in over 3,700 user sessions per day on average in the past two years. Human access through an easy-to-use graphical interface with browse and search capabilities, along with programmatic access via API with structured methods and data filtering, together allow for effective large-scale analyses. Our progressive automation of data collection, annotation, and release has accelerated the project growth by leveraging machine learning^29^ and modern cloud computing. This critical mass of public data and functionality promotes biomedical advances that are not possible without centralized coordination. We believe that the described NeuroMorpho.Org approach and methods may benefit other similar efforts and datasets in neuroscience and beyond.

## Materials and Methods

In the contemporary landscape of scientific data processing, the design of automation pipelines plays a pivotal role in enhancing efficiency and scalability. The NeuroMorpho.Org processing workflow epitomizes this approach through its innovative use of Dockerized ^24^ microservices. Each microservice functions as a modular building block that carries out a specific task or process within the overall pipeline, altogether referred to as a mesh grid architecture. This approach not only facilitates ease of updates and maintenance but also ensures that each component can be developed, tested, and deployed independently. The use of Docker containers for these microservices is particularly beneficial, offering a lightweight, portable, and consistent environment for each service regardless of the deployment platform. This encapsulation significantly reduces conflicts between services and streamlines installation across heterogeneous computing environments.

Thus, within NeuroMorpho.Org, one microservice is dedicated to the initial validation and ingestion of digital tracing files, ensuring that they conform to predefined schemas and standards ^28^. Other microservices perform feature extraction in the form of neural descriptors, such as persistence vectors^30^ and morphometric measurements^26^, from the standardized reconstructions. Those neural descriptors then serve as input to a similarity search microservice that flags potential duplicates upon data screening and allows users to find data morphologies of interest based on an archetype ^27^. This decomposition into dedicated services not only enhances the scalability of the pipeline but also allows for parallel implementation of new functionalities, thus accelerating both data processing and continuous development. The strategic progression towards a fully automated pipeline underscores a commitment to efficiency and reliability, with each microservice contributing to a seamless, cohesive data processing ecosystem.

The overall workflow consists of multiple steps including literature mining, file conversion and standardization, metadata annotation, and public data release. While we describe the outcome of this organization in the Results, here we provide essential technical details about the implementation.

The pipeline starts by identifying new peer-reviewed publications that contain neural reconstructions and actively contacting authors to request data submission. For this purpose, we have developed two open-source services. The first one is PaperBot, an automatic crawler that identifies potential publications from multiple publisher portals^31^ based on massively parallel weekly keyword queries. The second tool is LiterMate, a classifier that uses natural language processing and deep learning to assess the likelihood of a publication to describe new neural reconstructions with high accuracy^32^. Upon identification of the target publications, we invite the corresponding authors to share their dataset via NeuroMorpho.Org ^33^.

Once the authors upload their data to our secure file server, a second set of microservices comes into play. The first step requires converting the original files to the standardized SWC format^34^ using the recently released web and API service xyz2swc^28^. This software imports all known (50+) file formats generated by any commercial or non-commercial morphology tracing system utilized in the scientific literature, including Neurolucida^35^, Imaris (Oxford instruments), Amira (Thermo Fisher Scientific, USA), Simple Neurite Tracer (SNT^36^), NeuronJ^37^, Knossos^38^, NeuTube^39^, Vaa3D^40^, and Trees Toolbox^41^.

The next step entails the auto-processing of the SWC files, which identifies any potential irregularity within each reconstruction. If a digital tracing contains included side branches or overlapping points, custom-made python programs remove those programmatically. In addition, an AWK code parses files to insert missing somas, reassign structural domains (e.g., neurites or glial processes), rescale to physical units, and correct non-positive diameters. More complicated edits require the manual intervention of our well-trained team of curators under visual inspection. Clear mistakes in the reconstructions, such as disconnected branches, are corrected whenever possible, while less obvious issues are simply listed in log files, which are also made available with the original and processed data. To complete this step, python and JAVA scripts automatically generate PNG images and morphometric measurements^42^, respectively.

Following data processing, we annotate the metadata associated with the reconstructions leveraging a specialized web portal to ensure terminological consistency via controlled vocabularies^43^. Moreover, the metadata portal facilitates annotation with a novel suggestion system that relies on machine learning-based information extraction via natural language processing^29^.

### Agile release workflow

The public upload of data and metadata on the NeuroMorpho.Org web site now works on a dataset-by-dataset basis, also known as agile release. We have not fully described this process before, and we therefore do so here. This multi-stage procedure consists of three main phases: automated quality control, data ingestion for author review, and main site update. Each phase leverages automated checks with administrator approval to maintain the integrity, completeness, and uniqueness of the curated neuroanatomical data on the platform.

#### Automated Quality Control

The initial phase of the agile release workflow is initiated by a NeuroMorpho.Org administrator through a web-based user interface, in turn relying on a backend Application Programming Interface (API), in which all the subsequent actions can be called in sequence or individually. The phase begins with the retrieval of data from the file server hosting the original and processed neural tracings, logs of edits and residual changes, generated images and measurements, and metadata. A series of automated scripts evaluates the integrity, completeness, naming, and extensions of all files, matches every reconstruction with ancillary information (images, logs, and measurements) and corresponding metadata, and flags any missing component. When multiple neural cells are reconstructed from a single original file, our software creates a symbolic link structure, which reduces space utilization on the server and speeds up the following ingestion. In the absence of errors, an automated similarity search^27^ looks for potential duplicates between the new submissions and the existing entries using the combined features of summary morphometrics and persistence vectors. This step is crucial for maintaining the uniqueness of the data in the repository, as collaborating labs may accidentally resubmit the same reconstructions used in distinct publications. A senior curator then visually confirms or rejects duplicates through a user interface.

#### Data Ingestion for Author Review

Authors’ review of data and metadata prior to public release is a critical step of the of NeuroMorpho.Org commitment to transparency and quality control. This phase ensures that the authors can securely and effectively inspect their submissions as they will appear to the research community. The initial step of this phase transfers the neural reconstructions, along with their metadata, ancillary files, and associated symbolic link structures, to a password-protected website. This secure environment allows only authorized individuals, namely the contributing authors, to access their dataset for review. Following the transfer, the dataset is transactionally ingested into a relational database. This guarantees the successful completion of the operation or, in the event of an error, enables a seamless roll-back without leaving the database in an inconsistent state. The final step of this phase involves assigning a new review release version, which facilitates multiple correction requests by the contributing authors, if needed.

#### Main Site Update

The final phase of the workflow involves preparing and integrating the contributing author-vetted dataset into the public NeuroMorpho.Org site for free community access and use. First, the validated file structure is replicated from the review site to the separate main (production) site, ensuring consistency of the released dataset and transactional integrity relative to the latest version approved by the contributing authors. Next, the automated service assigns a new agile release version, updating the web server and database accordingly.

The search indexing is then regenerated, currently using Solr cores, to enhance the speed of subsequent web user interface and public API queries. Finally, the update is announced on X (formerly Twitter) as well as on the NeuroMorpho.Org homepage, informing the community of the latest addition.

## Results

Launched 18 years ago with just 932 neuronal reconstructions from 7 animal species and 23 labs, NeuroMorpho.Org recently passed 260,000 neuronal and glial (“neural”) reconstructions from more than 90 species - from C. elegans to human - and nearly 1000 labs worldwide. Unlike single-institution databases, these freely available data are truly diverse, with >1400 cell types, including long-range projection neurons, local interneurons, sensory receptor cells, a variety of glia, and 400 brain regions from both vertebrate and invertebrate nervous systems. Animal subjects include all common laboratory models (e.g., mouse, rat, fruit fly, zebrafish, and macaque) but also rarer specimens such as lion, leopard, tiger, giraffe, panda, elephant, bonobo, dolphin, whale, sloth, bat, anteater, and many more.

All NeuroMorpho.Org reconstructions are downloadable as SWC files^21,34^, a data format for which our team has led the community standardization following FAIR (Findable, Accessible, Interoperable, Reusable) principles^44,45^. In all cases, we publicly release both the original and processed data along with the log of all changes and residual notes (Figure 1, NMO_145047^46^). Every reconstruction is also accompanied by rich metadata. This information specifies details on the animal subject (e.g., species, strain, sex, developmental stage, age, and weight), anatomy (brain region and cell type), experimental protocol (labeling, objective type, and magnification, slicing direction and thickness), and provenance (contributing lab and bibliographic reference)^13^. We also augment all reconstructions with detailed morphometric measurements, such as total arbor length, number of branches, and fractal dimension^42,47^. The metadata and morphometrics enable both more powerful search and filtering capabilities, and stand-alone meta-analyses via conveniently retrievable summary reports^26,27^, adding functional value to the data.

**Figure 1.**
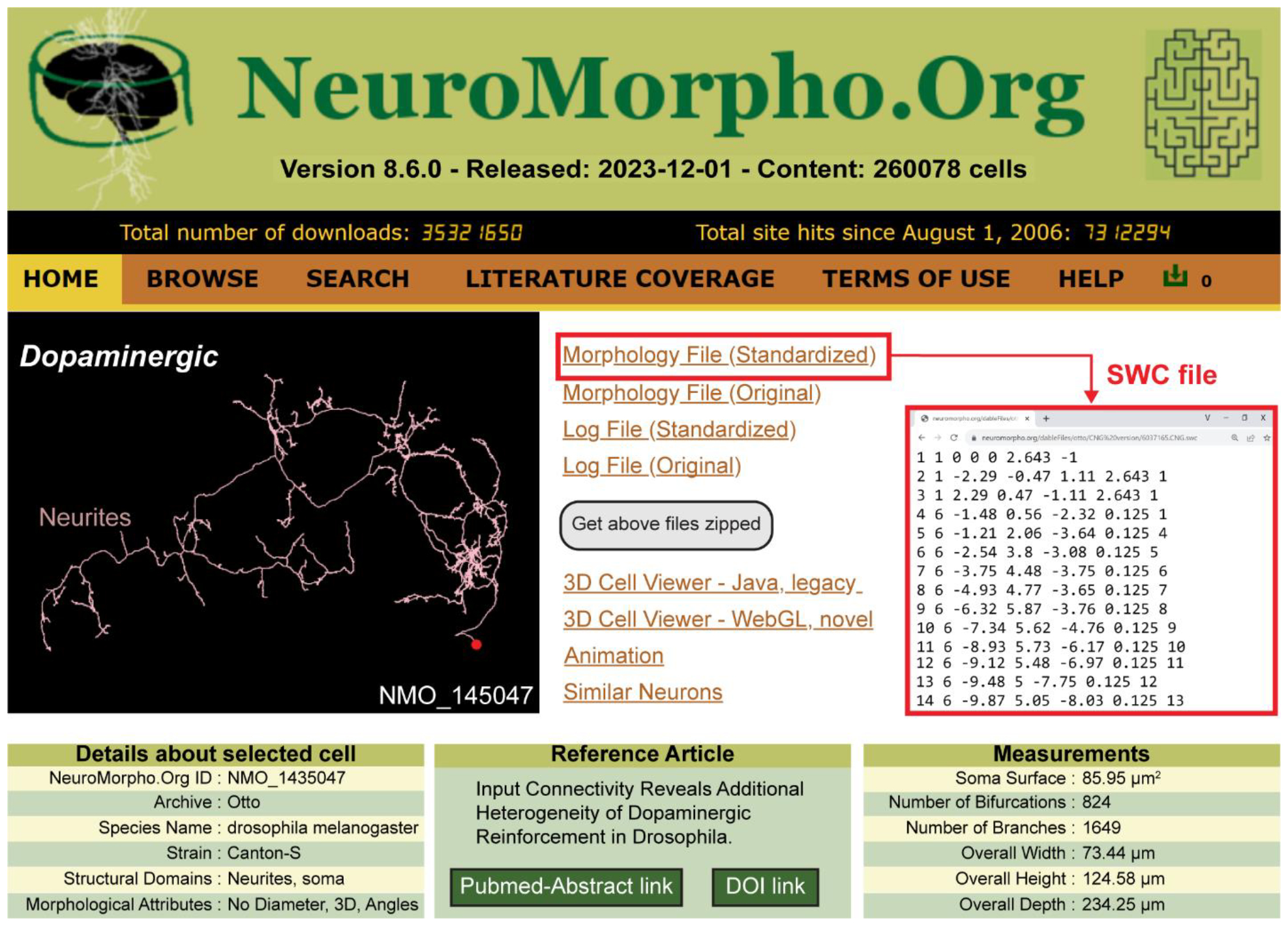
NeuroMorpho.Org is a public repository of neural morphology. **Top**. Version 8.6 (December 2023) contained over 260,000 digital reconstructions from nearly 1000 labs worldwide. **Middle left**. Representative reconstruction from a dopaminergic neuron (NMO_145047). **Middle right**. Links to original data, standardized files in SWC format (red box), 3D viewers, and similarity search. **Bottom**. Examples of metadata annotation fields, reference article, and morphometric measurements for the illustrated neuron.

The Terms of Use of NeuroMorpho.Org are prominently articulated on the website. Besides providing a liability disclaimer, they require users to mention NeuroMorpho.Org as the data source and to cite the original peer reviewed publication in which the downloaded data was first described. These requirements create a mechanism for author recognition and credit attribution. Moreover, they incentivize scientists to share data by ensuring a positive return on their impact metrics, such as h-index, via citation growth^48^. Data and tool usage is otherwise completely permissive along the most liberal Creative Common CC-BY and MIT licenses, respectively.

The analysis presented here is based on the content of NeuroMorpho.Org v.8.6.8 (March 2024), consisting of 1452 archives. Of these, the first 981 archives were released between 2006 and 2019 (v.1.0-v7.9), while 471 were released starting with v.8.0 in 2020 utilizing agile releases.

### Increase of NeuroMorpho.Org content

To properly analyze the remarkable growth of this database since inception, it is useful to consider the context of the overall data production in the field. The commercial technology to digitally reconstruct neural morphology was practically introduced in the early 1980s^49^, but became more popular at the turn of the millennium. Thus, when NeuroMorpho.Org was launched in 2006, we requested from the authors the relevant data published up to then, and subsequently proceeded to invite the deposition of new data as new publications continued to appear.

The total amount of neural reconstruction data described in the literature increased steeply over the past quarter century (Figure 2), from fewer than 2000 traced cells per year in 2000 to over 180,000 in 2023. Moreover, while initially a small fraction of the data produced was shared (<20% on average until 2008), the tide clearly turned more recently ^50^, with >80% of shared data in the last five years (Figure 2A). As a combined result of the overall growth of data and of the proportion of shared data, more than half of the neural cells ever reconstructed have been available at NeuroMorpho.Org since 2019.

**Figure 2.**
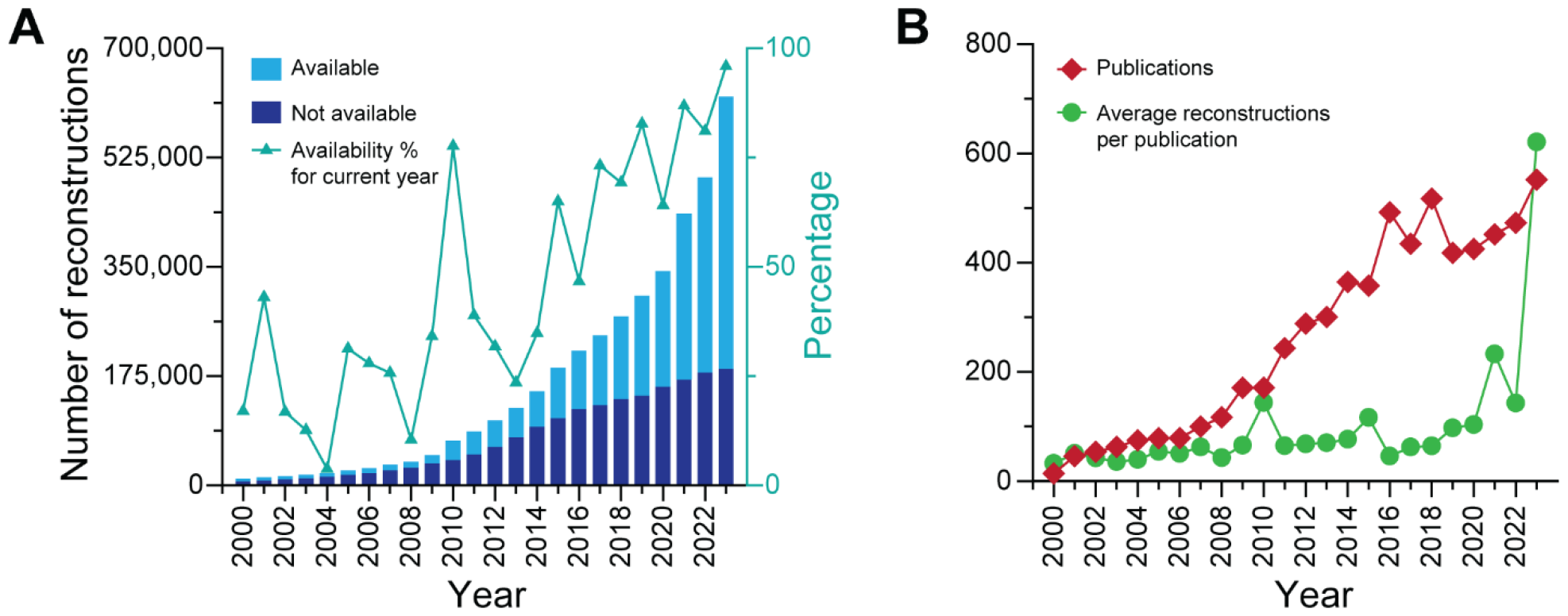
Growth of neural reconstruction data. **A**. Bar stacked plots of the cumulative numbers of available (light blue) and unavailable (dark blue) reconstructions by year. The turquoise line plot shows the corresponding percentage of available reconstructions for each year. **B**. Yearly numbers of publications describing reconstructions (red) and of reconstructions per average publication (green). All 1985-2000 publications are averaged together in the year 2000 data point.

It is also interesting to investigate whether the increase in the amount of neural reconstruction data is due to a greater number of publications in the field or to a tendency for authors to include, and for referees to expect, larger and larger dataset to support a published scientific claim. Our analysis shows that both factors are at play (Figure 2B). In particular, both the number of yearly publications describing morphological reconstructions and the average dataset size grew by an order of magnitude during the analyzed period, from fewer than 50 to more than 500.

### Which neural reconstructions are not available and why?

Next, we asked whether the willingness to share reconstructions differs across distinct research communities (Figure 3). To address this question, we analyzed the amount of available and unavailable data among the most common animal species, cell types, brain regions, and tracing systems (Figure 3A). Approximately half of the rodent tracings are available at NeuroMorpho.Org (48% for rats and 58% for mouse); this proportion is higher for humans (66%) and Drosophila (88%). Clear differences are also noticeable between the proportion of shared neurons (52.2%) and glia (82.9%). These results can be readily understood in light of the previously described trend of increasing availability over time. In the early days of digital reconstruction most neuroscientists largely focused on rat (and then mouse) neurons, and during this period only a minority of the data were shared. More recently, research emphasis expanded to encompass the fruit fly model ^51^ and technology advanced to enable human ^52^ reconstructions. Meanwhile, the important physiological and pathological role of non-neuronal cells, such as microglia, astrocytes, and oligodendrocytes, became increasingly recognized. By the time glia (and drosophila) tracings rose to prominence in peer-reviewed publications, most authors were willing to share their data. This interpretation is consistent with both the relatively uniform proportion of sharing across brain regions, and the greater availability of reconstructions collected with relatively more recent tracing systems, such as SNT and Imaris, relative to more established earlier systems like Neurolucida and NeuronJ.

**Figure 3.**
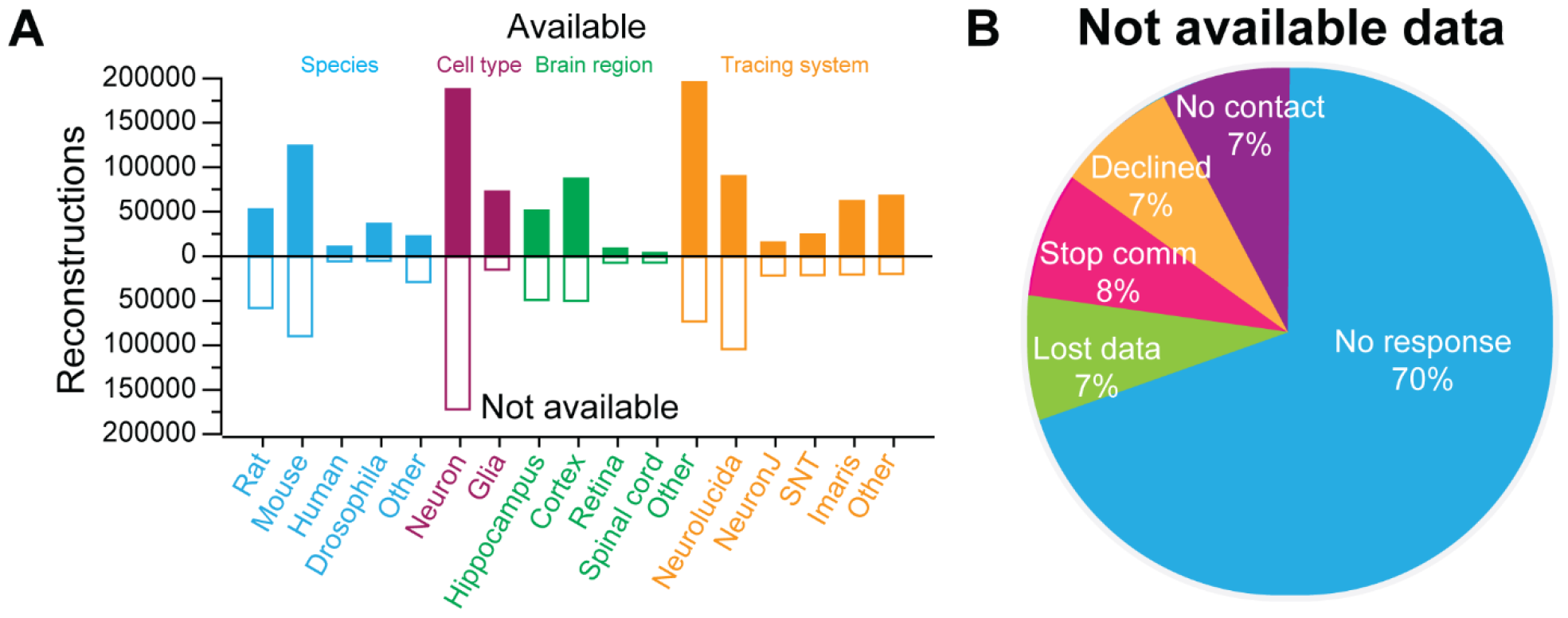
Breakdown of reconstruction availability. **A**. Number of available (filled up bars) and unavailable (hollow down bars) reconstructions by selected species (blue), cell types (purple), brain regions (green), and tracing systems (yellow). **B**. Main reasons for data unavailability (Stop Comm: authors stopped communication after an initial positive response).

We then analyzed the main reasons for not sharing data ^53^. The principal culprit appears to be lack of time or commitment, as the absolute majority of authors not sharing reconstructions simply fail to respond to the collegial invitation and subsequent reminders (70%). An additional 8% among the invited authors not sharing data stop communicating after an initial positive response, while only 7% flatly decline to share their data. The remaining 14% of authors not sharing are equally divided between those who declare the reconstruction data no longer available (due to computer crashes, moving institutions, or personnel change) and those with no valid email contact due to retirement, leaving academia, or decease. It is important to reiterate that these results only pertain to the (now minority) of reconstruction data that are not available, while most authors in recent years choose to share their data via NeuroMorpho.Org.

### Status of NeuroMorpho.Org processing pipeline automation

NeuroMorpho.Org data sharing is a multifarious effort involving request, standardization, and annotation of reconstructions and metadata into a community database. Our recent automation of this process has enabled increased data sharing capacity and community engagement for novel applications independent of the intended purpose of the original data collection. The pipeline, now largely automated by microservices, is divided into sequential steps, namely literature mining, metadata annotation, and data ingestion for public release (Figure 4).

**Figure 4.**
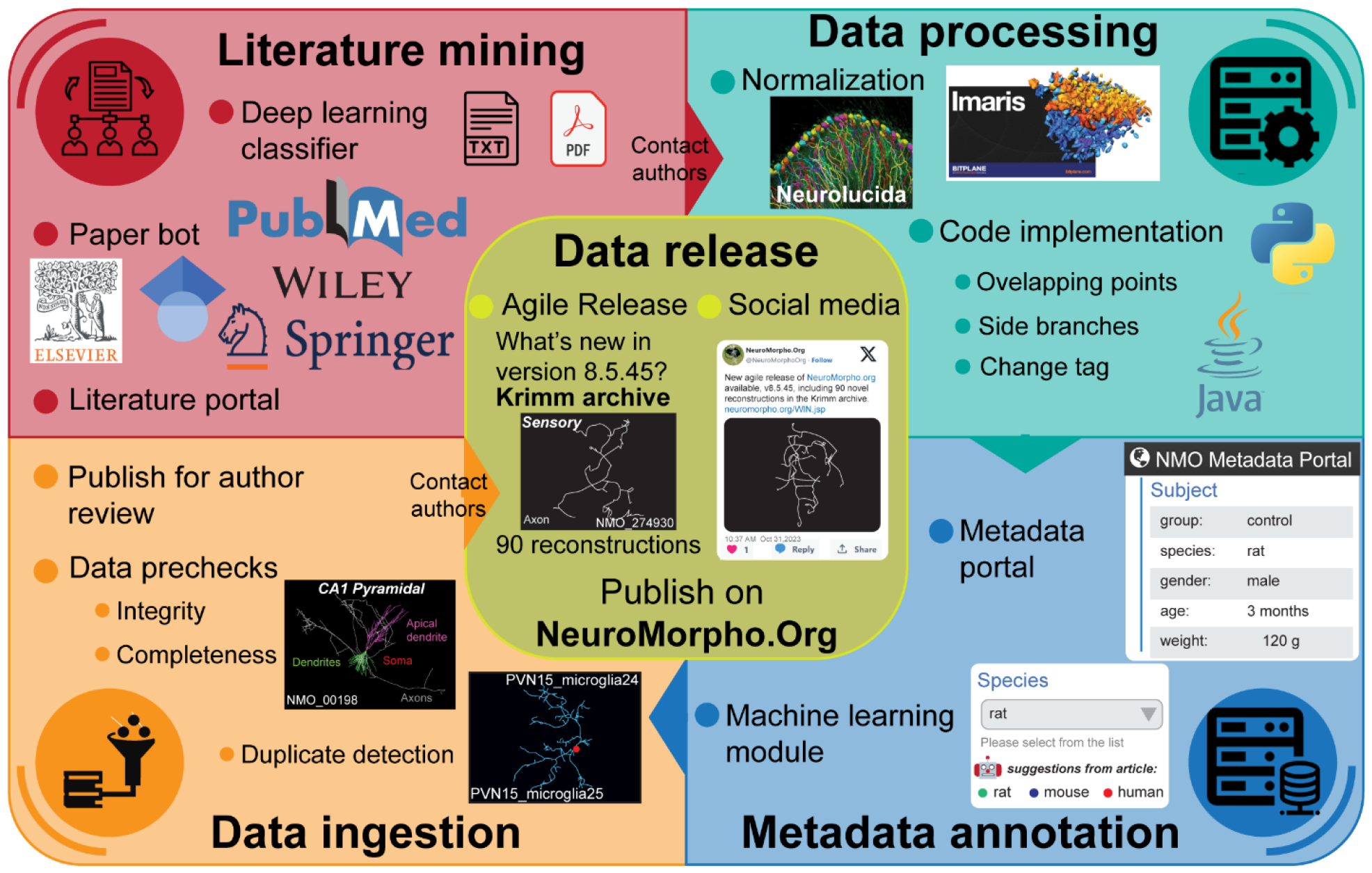
NeuroMorpho.Org processing pipeline. **Literature mining** harnesses automated tools for searching possible publications associated with neural reconstructions (PaperBot) and a deep learning classifier to help identify the relevant articles (LiterMate). **Data processing** starts by converting all tracing data from multiple file formats to SWC, followed by resolving common irregularities in the reconstruction data. **Metadata annotation** leverages a web-based portal with a backend machine learning module that now automatically suggests the most likely entries extracted from the original publication. **Data ingestion** includes verifying the integrity, uniqueness, and completeness of each reconstruction, and an author review stage prior to public release. **Data Release** follows an agile (dataset-by-dataset) release strategy accompanied by social media posting to advertise the newly added reconstructions.

#### Literature mining

We have over the years implemented highly optimized literature searches to identify every scientific publication describing digital reconstructions of neural morphology. Our algorithms employ deep learning to autonomously mine all publisher portals continuously and in parallel. This allows us to invite authors to deposit their datasets within days of the publication date, while they still supposedly have convenient access to the data. This increases the probability of a positive response to the data sharing invitation, given that the authors have recently organized the data for analysis and the trainees are still in the lab. An increasing proportion of authors now spontaneously contact us upon manuscript submission to deposit their neural reconstructions at NeuroMorpho.Org prior to publication acceptance. Only publications that are evaluated as low confidence by the automated service remain to be evaluated by our curation team, reducing human labor by 80%.

#### Data processing

We subject all reconstructions provided by the authors to a consistent workflow of conversion, standardization, and quality control. This pipeline is now largely computerized, but each outcome is visually proof checked by trained curators. Only reconstructions that require specialized handling are processed manually. The developed suite of custom-made software programs reduced the average processing time per reconstruction from days to minutes.

#### Metadata annotation

To enable easy annotation, the NeuroMorpho.Org metadata portal guides the authors (or alternatively our curation team) through a series of proposed pre-annotated entries for each metadata field (for example, “species: mouse”). These entries are automatically pre-extracted from the full text of the corresponding peer-reviewed publication. If deemed correct, the user can accept an entry with a simple click, or by simply skipping to the next metadata field. If warranting a correction, a convenient dropdown menu includes all relevant terms for that field (e.g., all species names) that already exist in the database, with the added features of autocompletion. Only when not finding the desired term, the web form provides a free-text option. All new terms are double-checked by our curation team and listed explicitly on the main site for community feedback at major releases at least once per year. Adoption of the metadata portal service more than halved the annotation time relative to the previous spreadsheet-based method.

#### Data ingestion for public release

After a series of prechecks to ensure the integrity, uniqueness, and completeness of each reconstruction, we invite authors to preview their data and metadata on a password-protected review site, allowing them to request any desired change prior to public roll-out. Finally, the dataset is released publicly with two simultaneous community announcements, each indicating the new agile release version, the name of the archive, and the total number of added reconstructions. The first announcement is posted directly on the NeuroMorpho.Org website, under the “What’s new” section. The second announcement is tweeted via social media on X (and embedded on a web feed on the NeuroMorpho.Org home page).

### Average processing time per neural reconstruction

To assess the impact of the incremental automation on the overall processing time, we quantitatively analyzed the project records since the inception of the project. Back in 2005, not even a single reconstruction was processed per day; during the first 3 years after the official launch, 4.3 ± 2.8 reconstructions were processed per day on average. During 2017-2019, when the glial cells were formally added to the database, the average number of daily processed reconstructions increased to 54.1 ± 4.9. In the last three years, this value increased further to 199.1 ± 111 per day. In summary, we passed from processing one tenth of a reconstruction per days to almost one reconstruction per minute (Figure 5A). A complementary way to gauge the effects of automation is by clocking the average processing time per reconstruction. This value dropped from 9.6 hours during the first years of the project to 3.5 ± 1.2 minutes in the last three years (Figure 5B).

**Figure 5.**
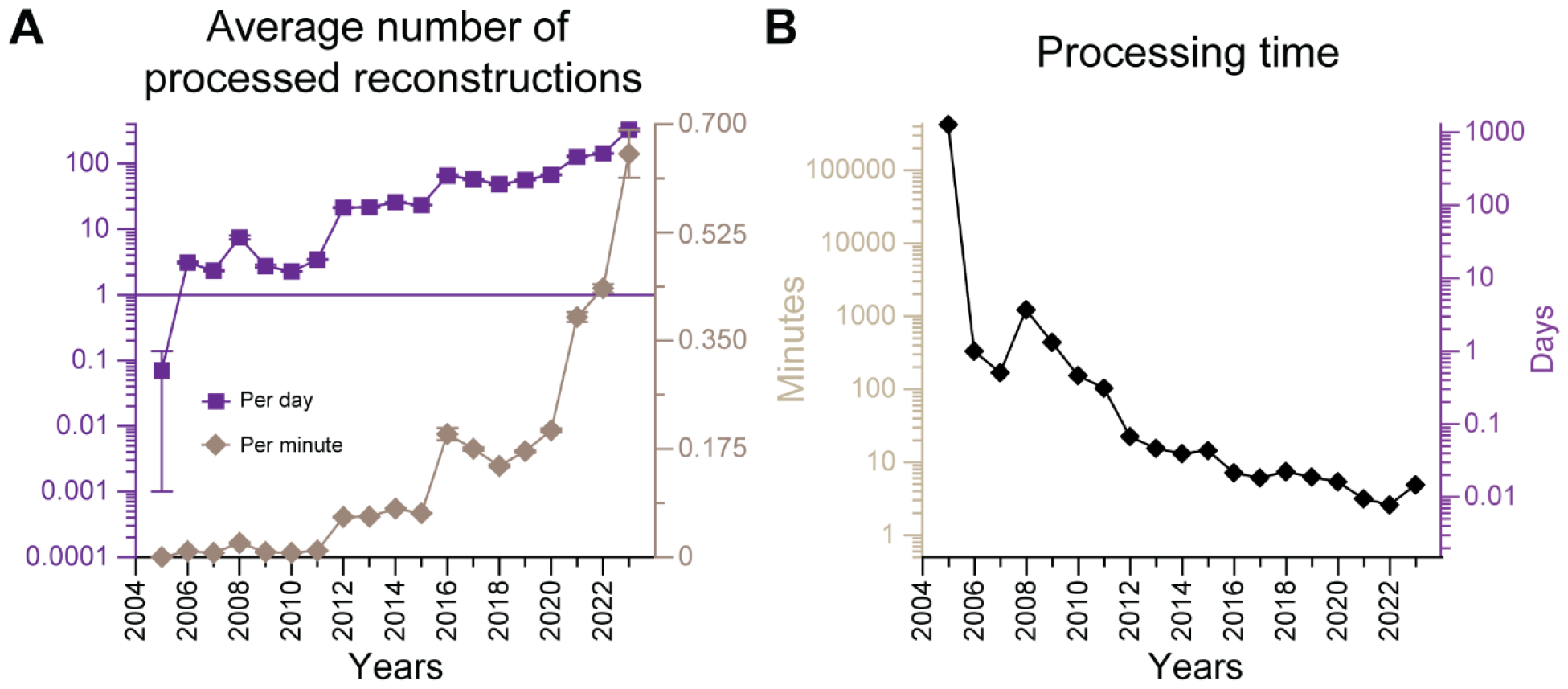
Processing throughput of neural reconstruction over the project lifetime. **A**. Average number of processed reconstructions per unit of time in days (purple) or minutes (brown) **B**. Average time to process one reconstruction in days (right axis) and in minutes (left axis).

### Commonly required edits during processing

Next, we evaluated the number and types of manual processing interventions required to release the data, if any. This analysis includes the (most recent) 574 archives (out of 1452) for which we kept the relevant records. Almost half (46%) of these archives were fully completed by automated processing, usually resulting in public release within less than a week from data receipt. A similar proportion of archives involved one or two edits (22.5% and 22.1%, respectively), while fewer than 10% of archives required three or more edits (Figure 6A). The manual intervention time for each edit ranged from minutes to hours depending on the complexity of the task. The most commonly required edit consisted of repairing long connections (15.7%), followed by separating cells jointly traced within the same file (9.5%). Less frequent edits included soma correction (3.4%), removal of non-neural structures like anatomical contours (1.6%), and miscellaneous others (altogether accounting for 12% of cases), such as visually identifying and tagging distinct structural domains, for instance apical vs. basal dendrites (Figure 6A). We then analyzed the association between specific types of edits and individual tracing systems (Figure 6B). The most notable examples are separating multiple reconstructions from single files most commonly occurring in Imaris (25%), deleting doubly traced branches, often encountered in Simple Neurite Tracer (10.6 %), and removing contours, almost exclusively required in Neurolucida (4%). While the time needed to manually separate reconstructions from single files depends almost linearly on the number of neural cells in question (up to several hours for hundreds of cells per file), removal of contours is usually a matter of minutes in most cases.

**Figure 6.**
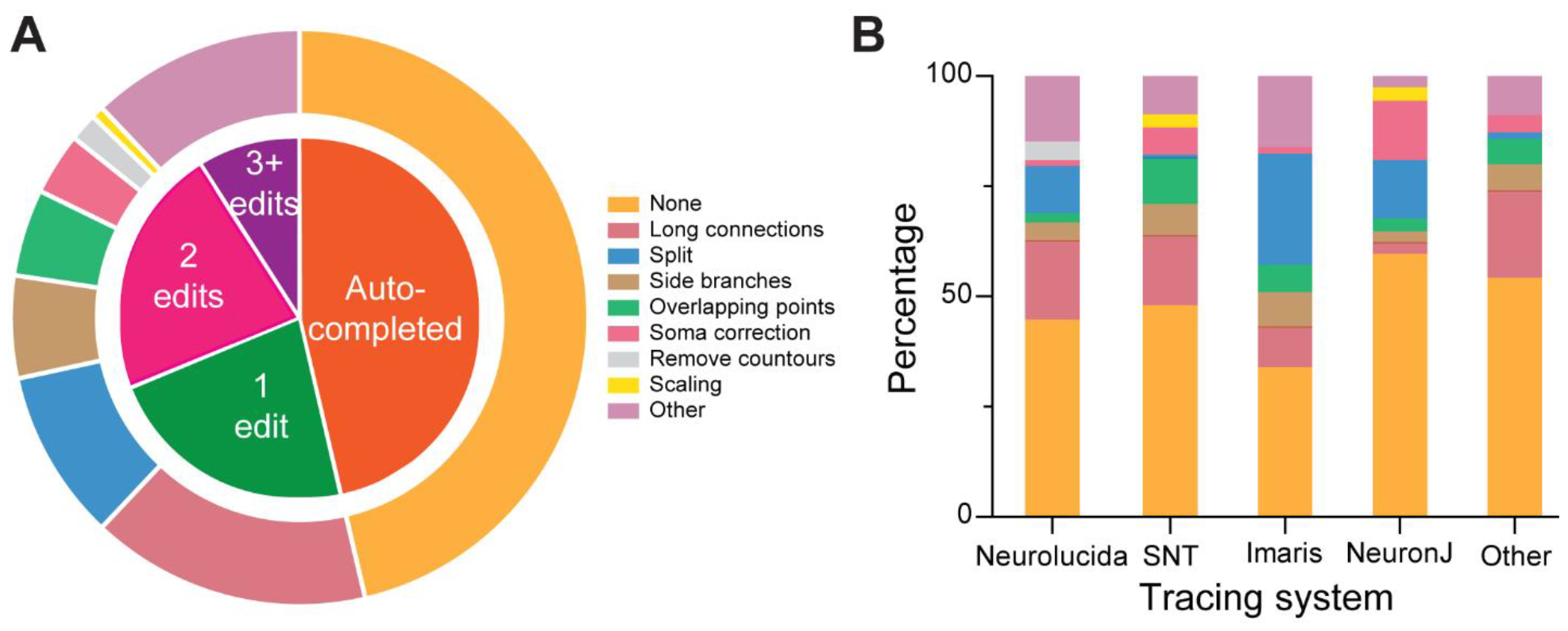
Common data edits during processing. **A**. Number (inner pie chart) and types (outer pie chart) of manual data processing interventions required prior to release, if any. **B**. Proportion of edits associated with the most popular tracing systems.

### Geographical distribution of NeuroMorpho.Org users

NeuroMorpho.Org is engineered to ensure the broadest worldwide dissemination of laboriously acquired experimental data and associated metadata. The fully documented API and intuitive graphical user interface enable advanced querying capabilities that provide unhindered access to the data and all accompanying information. The end result is an extraordinarily wide distribution of users, with confirmed downloads from 210 out of the 253 countries and territories in the world (Figure 7). Access to this resource has proven to be particularly impactful for emerging research nations with limited or no funding for tracing neural cells by enabling creative data reuse through a standard internet connection. Notably, 78 % of the digital reconstructions currently included in the database were generated in North America and the European Union (EU), and none in Africa. Only 13.6 % of the laboratories that contributed data to NeuroMorpho.Org were located in the 152 countries defined by the International Monetary Fund (IMF) as “developing”, and none of these were in the 47 “least developed” (per IMF) countries.

**Figure 7.**
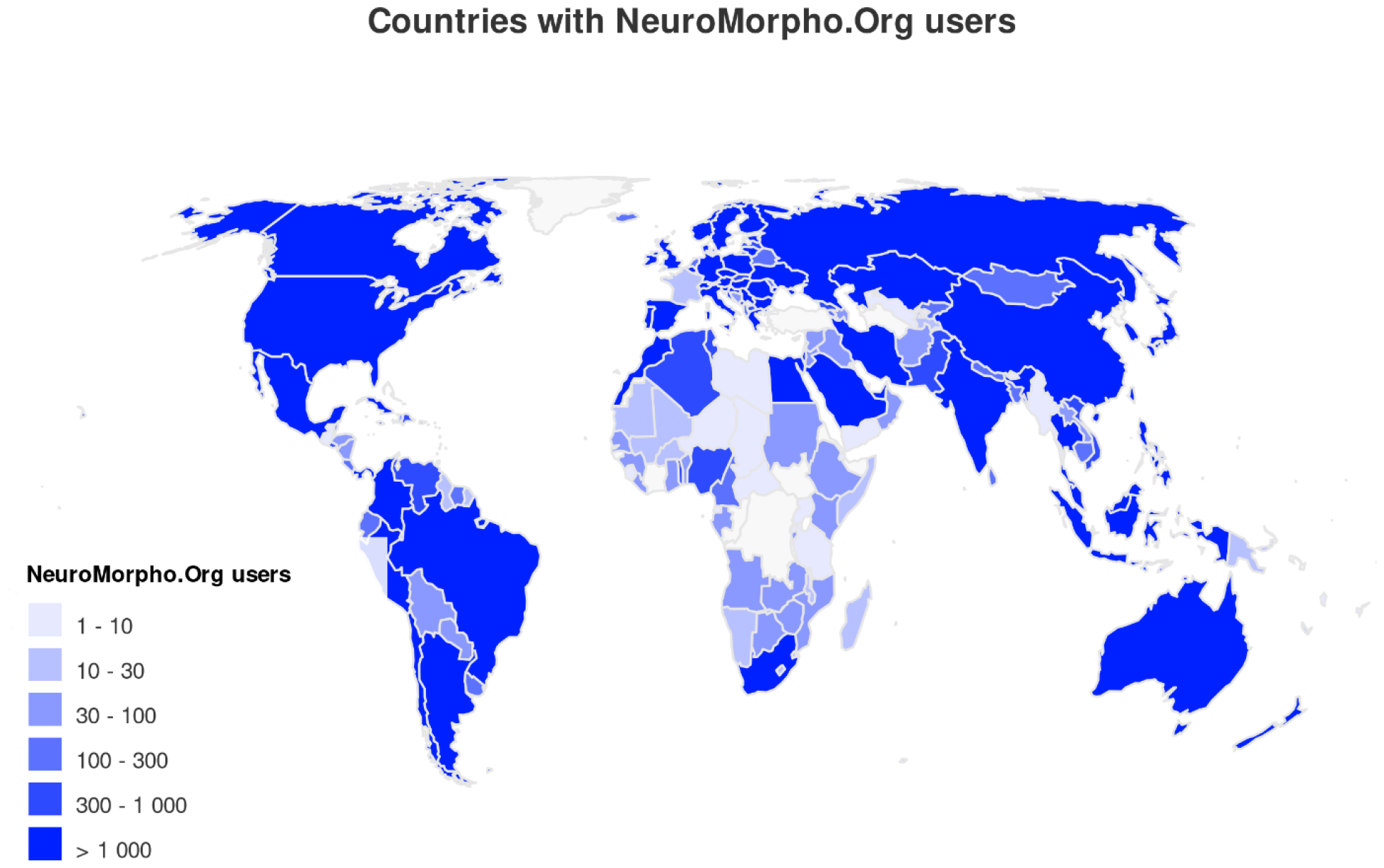
Number of NeuroMorpho.Org users by country since project inception.

Comparatively, only 87.6 % of all user sessions were from Europe and North America, and the remaining 12.4 % include almost all African countries. More broadly, 26.5% of the unique IP addresses accessing and retrieving data from NeuroMorpho.Org were located in IMF developing countries, and 0.9% of these were from the IMF least developed countries (these latter ones tallying as many as 450 unique IP addresses). As an additional single example, Greece has no contributing labs, but constitutes a major user base of NeuroMorpho.Org, with 0.2 % of all user sessions. This usage from emerging research nations has grown over the years, demonstrating increasingly pervasive penetration of the database throughout the broader international community.

More than 32 million downloads from NeuroMorpho.Org have already yielded >800 new publications by the research community in journals such as Nature, Science, Cell, New England Journal of Medicine, Trends in Neuroscience, Nature Neuroscience, Neuron, Cerebral Cortex, Journal of Neuroscience, Nature Communications, PLoS Biology, eLife, Current Biology, and Nature Methods, as well as preprints like BioRxiv and dozens of doctoral dissertations. Scientific findings and applications span the gamut of meta-analyses, computer simulations, comparative morphology, data mining, hypothesis-testing, and algorithm design. These discoveries were made possible not only by sharing data, but also by the sheer size and diversity of the shared data, allowing inferences otherwise beyond the reach of individual labs. For example, over 1300 experimental conditions are represented in NeuroMorpho.Org, including 623 genetic manipulations, 242 pharmacological treatments, 63 surgical lesions, 63 behavioral interventions, and 357 combinations thereof.

## Discussion

NeuroMorpho.Org has supported research progress in neuroscience for over 17 years. During this time, scientists who have interacted with the database and the SWC format as junior trainees are now continuing in their efforts as senior researchers. For thousands of practitioners, NeuroMorpho.Org is the go-to trusted resource for finding neural reconstructions and for depositing tracing data. NeuroMorpho.Org also provides immediate value to the broader community with 1-click accessible 2D, 3D, and interactive visualizations of all published neurons and glia. Consequently, in addition to extensive scientific usage, NeuroMorpho.Org is employed as an educational resource in high schools, colleges, graduate programs, online labs, and web courses^54,55^, and has inspired numerous multimedia artistic renderings including illustrations, sculptures, virtual reality displays, and audiovisual exhibits. In the private sector, DRVision Tech developed and launched their best-selling image analysis product, AIVIA5, using NeuroMorpho.Org for algorithm training.

Database content and web portal functionality are clearly documented in a QuickStart guide and Frequently Asked Questions. Furthermore, all our data processing tools, and standardization pipeline are available open source via GitHub ^56^. This not only allows full replicability, but also catalyzes a rich community ecosystem of inter-related resources providing complementary and synergistic opportunities for further development ^21^. In fact, a growing number of other resources replicate all or parts of the NeuroMorpho.Org content in new applications to accelerate scientific discovery. These are developments that we welcome, encourage, and facilitate. Examples include the Blue Brain Project’s Hippocampus Hub^57^, which is continuously synched with relevant NeuroMorpho.Org content; multiple informatics pipelines enabling interactive queries, analyses, and simulations directly from NeuroMorpho.Org; and an external service sourcing NeuroMorpho.Org data to print 3D neuron models.

Thus, in line with CARE principles^58^, NeuroMorpho.Org maximizes the collective benefit of research, regardless of affiliation, nationality, gender, belief, or ethnicity. For scientists providing the original neural tracings, participating in NeuroMorpho.Org reduces the time and labor required to address each inquiry about their data, while increasing recognition and impact of their research. We provide clear and simple instructions for how to upload datasets, and the association of each reconstruction with the peer-reviewed publication in which it was originally described ensures authority control. From the user’s perspective, our endeavor removes all barriers to data access. The annotation of every downloadable file with detailed metadata expands the capability and capacity of the original data while allowing users to make informed decisions about responsible scientific applications. From the layperson standpoint, reuse of research data multiplies the profit of scientific investments and reduces the cost of animal subject lives.

Hence, the organization of NeuroMorpho.Org supports the ethical pillars of maximizing benefits, minimizing harm, promoting equality, and preserving future opportunity for non-exclusive reuse. The overall scientific impact of NeuroMorpho.Org is summarized by nearly 4000 peer-reviewed publications: 2238 describing data available through the database, 844 using downloaded reconstructions, 53 publications about the project itself, and 865 additional references citing NeuroMorpho.Org, often as an exemplary resource in neuroscience data sharing. Noteworthy breakthroughs with immediate and long-term clinical implications enabled by NeuroMorpho.Org include optimizing placement of intracranial electrodes in human neurosurgery ^59^ and predicting neurological radiation damage of cancer radiotherapy or long-haul space travel ^60^. Much additional progress resulted from reuse of data shared by NeuroMorpho.Org, such as linking diffusion anisotropy with neurite development ^61^; identifying systematic differences of cell-type specific branching geometry in the cortex of mice, monkeys, and humans ^62^; inferring communication signals in visual motion detection ^63^; revealing the mechanisms of circuit wiring ^64^; deconstructing the contributions of voltage-gated and ligand-gated channels to synaptic integration ^65^; correcting staining method biases in neuroanatomy ^66^; demonstrating Pareto optimality between cable minimization and conduction delay ^67^; and quantifying systematic patterns of structured connectivity ^64^.

If peer-reviewed publications provide an index of retrospective utilization, grant applications and awards typically reflect prospective usage. The National Institutes of Health (NIH) has received at least 366 proposals mentioning NeuroMorpho in key sections (titles, specific aims, and summary statements). The applications came from 255 Principal Investigators and 124 external organizations. As many as 142 of these proposals have already been funded to date, with $246M projected to be awarded and spent.

Essential to NeuroMorpho.Org is its commitment to both open-source software development and open-standard formats. The goal is to maximize data production and distribution while minimizing cost. Until four years ago, 81% of digitally traced neural morphologies utilized commercial technology whose high cost (∼$30,000 per workstation license) constituted a steep entry barrier for many smaller labs. Moreover, the several dozens of proprietary file formats for neural reconstructions, each accounting for less than 10% of the data, created a Babel Tower scenario hampering exchange and cooperation. We catalyzed the design and implementation of new freeware tracing algorithms by organizing community challenges (DIADEM^68^) and consortium activities (NIH mini-symposium at Society for Neuroscience), releasing gold standards, and sponsoring hackathons (BigNeuron^69^). As a result, 45% of all neural morphologies reconstructed in the past four years used free digital tracing software. Furthermore, we collaborated with community stakeholders to produce, integrate, and distribute open-source software for converting all known file formats for digital morphology into the SWC lingua franca, nurturing this nascent community standard into practical coalescence. The outcome is a tremendous increase in both data production and cooperative exchange among data users and producers.

Our standardization efforts precipitated wider adoption by the research community. Many tool providers now support the SWC standard format, including NEURON ^70^, the popular software for physiological modeling, as well as GTree ^71^ and Fast Tracer ^72^ algorithms, which automate reconstructions of neural arbors from microscopy image stacks, and even corporate enterprises like MicroBrightField. Therefore, our efforts had both a “downstream” impact, by streamlining data import into analytics tools, and an “upstream” impact, by standardizing export of neuronal reconstructions into a common format. Since inception, NeuroMorpho.Org has continuously matured in resonance with the evolving scientific interests of biomedical research. For example, mounting community recognition has focused attention on the foundational physiological relevance of glia in brain function and devastating diseases such as epilepsy, senile dementia, and stroke. Notably, NeuroMorpho.Org inspired other biomedical research communities to adopt the SWC standard for their data, such as to reconstruct cerebral vasculature from Magnetic Resonance Angiography, a crucial diagnostic exam for detecting brain aneurisms. Recently, the non-profit International Neuroinformatics Coordinating Facility (INCF) ^44^ officially endorsed SWC as an INCF standard.

NeuroMorpho.Org has transformed collaborative data sharing from rare peer-to-peer occurrence in cellular neuroanatomy to a valuable mainstream activity. Consequently, two-thirds of the digital neural morphologies ever reconstructed since the dawn of computer-interfaced microscopes (1982) are now publicly available at NeuroMorpho.Org. We have pioneered a strategy of transparent online publication of all our data sharing requests and of their outcomes to promote community awareness. This approach resulted in a remarkably broad base of labs willing to share their data: 78% of all tracings received to date were uploaded by 1st-time contributors. In fact, incentivizing career scientists to share their data is central in our design. We have implemented novel algorithms to identify every re-usage of individual datasets in peer reviewed publication, thus quantifying the ‘added value’ of each contributing lab^48^. Authors can use this evidence-based information for career promotions, merit reviews, award nominations, and grant renewals. We also store the grant support of each dataset submission, enabling funding agencies to track the impact of their investments by download and re-use in addition to standard publication counts. The long-term goal of the project remains to always foster public availability of data to the maximum possible extent. Further ongoing automation may soon allow further community involvement, such as one-click data submission and distributed wiki-curation of published content. NeuroMorpho.Org demonstrated that broad adoption and responsible stewardship are the strongest insurances for long-term sustainability and durable biomedical impact.

## Acknowledgments

We thank the entire NeuroMorpho.Org team, past and current members, with special mentions to Cintia A. Martinez Cardenas, Valerie T. Nguyen, Drew Di Donna, Joseph Ndiaye, Lima Ghairatmal, Hafsa Rehman, and Sadaf Alizai, who presented partial preliminary results of an earlier version of this data analysis. We are also grateful to our software developers Lin Shen and Zengxin Li for their help and continuous support of our website. Finally, we would like to thank the Federation of American Societies for Experimental Biology (FASEB) and the Office of Data Science Strategy at the National Institutes of Health (NIH) for the Distinguished Achievement Award from the DataWorks! Prize to our team. This work was supported in part by grants R01NS39600, RF1MH128693, and R01NS86082 from the NIH. The funding sources were not involved in study design, data collection and interpretation, or the decision to submit the work for publication.

## Author Contributions

CT contributed to the conceptualization, methodology, investigation, data curation, formal analysis, writing original draft, visualization, supervision, and project administration. BL contributed to the conceptualization, methodology, investigation, writing original draft, software, visualization. GAA contributed to the conceptualization, methodology, resources, investigation, data curation, visualization, supervision, project administration, and funding acquisition. All authors were involved in revising and editing the manuscript.

## Data Availability Statement

The data that support the findings of this study are available in the methods and/or supplementary material of this article. These data were derived from the following resources at [https://NeuroMorpho.Org/,^73^].

## Footnotes

a When possible, relevant citations refer to publications describing morphological data publicly available at NeuroMorpho.Org.

